# Through-grid wicking enables high-speed cryoEM specimen preparation

**DOI:** 10.1101/2020.05.03.075366

**Authors:** Yong Zi Tan, John L. Rubinstein

## Abstract

Blotting times for conventional cryoEM specimen preparation complicate time-resolved studies and lead to some specimens adopting preferred orientations or denaturing at the air-water interface. We show that solution sprayed onto one side of a holey cryoEM grid can be wicked through the grid by a glass fiber filter held against the opposite side, often called the ‘back’ of the grid, producing a film suitable for vitrification. This process can be completed in tens of milliseconds. We combined ultrasonic specimen application and through-grid wicking in a high-speed specimen preparation device that we name ‘Back-it-up’, or BIU. The high liquid-absorption capacity of the glass fiber compared to self-wicking grids appears to make the method relatively insensitive to the amount of sample applied. Consequently, through-grid wicking produces large areas of ice suitable for cryoEM for both soluble and detergent-solubilized protein complexes. The device’s speed increases the number of views for a specimen that suffers from preferred orientations.

## Introduction

Advances in single particle electron cryomicroscopy (cryoEM) now allow routine high-resolution structure determination for many macromolecules and molecular assemblies. Nonetheless, the ability to prepare specimens suitable for imaging remains one of the main bottlenecks for these structural studies. Standard specimen preparation methods (Dubochet et al., 1988; Knapek and Dubochet, 1980) involve using a micropipette or syringe to transfer microliter volumes of solution onto the surface of an EM grid coated with a holey carbon or gold film (Figure 1*A*). Excess sample is removed by blotting with paper for seconds to produce a thin aqueous film, which is subsequently vitrified by plunging the grid into a cryogen such as liquid ethane or a mixture of liquid ethane and liquid propane. The liquid-absorption capacity of the paper is far larger than the amount of sample applied, suggesting that blotting for too long should lead to complete drying of the grid. However, once formed, these thin films of solution appear to be moderately stable, with continued blotting only slowly thinning the film (Armstrong et al., 2020). This continued thinning could occur either through further wicking or through evaporation.

**Figure 1.**
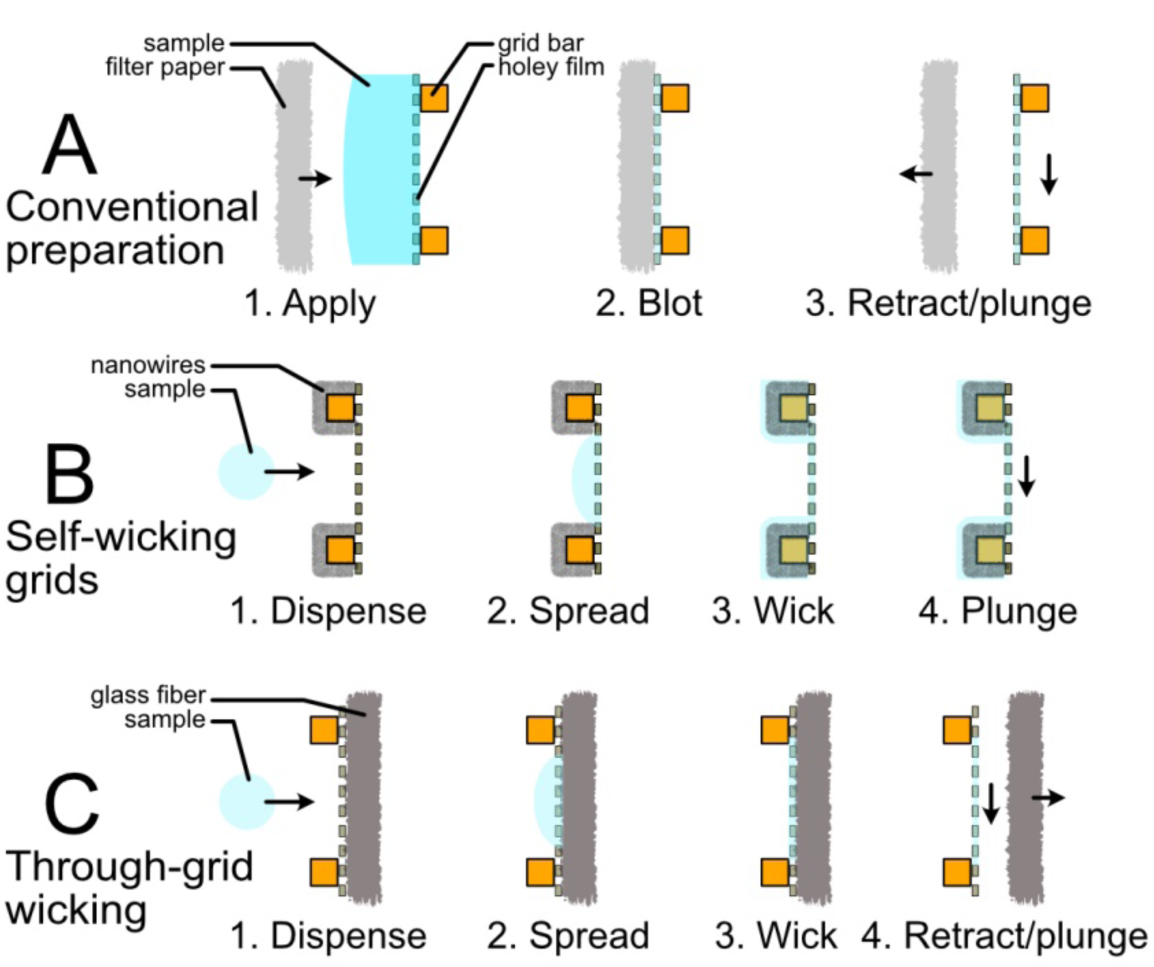
Cartoon of different wicking approaches for cryoEM specimen preparation approaches. (*A*) Conventional specimen preparation: the sample is applied to the grid and blotted with filter paper to produce a thin film suitable for plunge freezing. (*B*) Self-wicking grids: nanowires on the grid absorb excess liquid to produce a thin film. Small volumes of the sample are applied by a piezoelectric inkjet dispenser (Spotiton) or piezoelectric transducer (Shake-it-off). (*C*) Through-grid wicking: the sample is applied to one side of the grid while a filter on the other side removes excess sample to produce the film that will be vitrified.

Blotting with paper can be done from one or both sides of the grid. If blotting is performed from only one side, it is usually done on the side where the sample was applied. ‘Backside blotting’, where sample is applied on one side and blotting is performed on the other, has been used to concentrate filaments or particles that cannot pass through the grid holes (Amos and Hirose, 2007). Blotting a 3 µL sample into a 1000 Å thick film that covers a 3 mm diameter EM grid requires removal of 99.998% of the liquid. Consequently, blotting to remove sample without concentrating the molecule of interest on the grid is extraordinarily wasteful. The time required to prepare a thin film by blotting, which is likely due to the large volume of liquid that must be removed, prevents or complicates using grid freezing to study time dependent processes that occur on timescales shorter than a few seconds. During grid preparation, the sample exists as a thin liquid film with a large surface area to volume ratio, exposing fragile macromolecules to nearby hydrophobic air-water interfaces for an extended period. The interaction of protein particles with an air-water interface has been found to induce preferred orientations for many specimens (Noble et al., 2018) and cause denaturation of others (D’Imprima et al., 2019; Glaeser et al., 2016).

The introduction of self-wicking grids (Razinkov et al., 2016; Wei et al., 2018) provided a new way of preparing cryoEM specimens (Figure 1*B*). With these specimen supports, copper hydroxide nanowires are grown from the copper surface of copper-rhodium grid bars by incubating the grids in a solution of ammonium persulfate at basic pH. The grids are subsequently covered with a sacrificial film of holey plastic (Marr et al., 2014; Razinkov et al., 2016; Wei et al., 2018) that can be coated with carbon or gold before the plastic is dissolved. During specimen freezing, small volumes of liquid are applied onto the nanowire side of the grid. When the liquid contacts the nanowires, most of the sample is wicked to the grid bars, leaving a thin film suitable for vitrification and subsequent imaging. The volume of liquid applied is small and the wicking process occurs quickly, producing a suitable film in tens of milliseconds. However, the nanowires on the grid have a limited capacity to absorb liquid and can be easily saturated if too much sample is applied. Further, the speed of wicking from copper hydroxide-nanowire grids can vary from grid to grid, a problem thought to be due to variability in the copper used to make the grids (Wei et al., 2018). The small droplets of liquid needed for this approach have been generated previously by two different methods, although presumably other liquid dispensing techniques could also be used. In the first application of self-wicking grids, a piezoelectric transducer coupled to a glass capillary, similar to the printing head in an inkjet printer, was used to apply sample (Dandey et al., 2018; Jain et al., 2012). This device, named “Spotiton”, led to the commercial “Chameleon” instrument from SPT Labtech Ltd (Darrow et al., 2019). With Spotiton and Chameleon, the sample is applied onto a strip of grid squares as the grid is moved past the dispensing head, allowing the wicking behavior of the grid to be characterized with high-speed cameras. Suitable conditions are selected and the sample is applied to a different strip of grid squares on the same grid before plunge-freezing in cryogen. This approach requires high-precision liquid dispensing and high-quality optical components. Alternatively, liquid can be dispensed in a less controlled manner by a simple piezoelectric transducer like the ones found in ultrasonic humidifiers, an approach implemented with the open-source “Shake-it-off” (SIO) device (Rubinstein et al., 2019). This device relies on the piezoelectric transducer producing a distribution of droplets that varies in intensity across the grid so that some regions are saturated with liquid, some are dry, and some receive an optimal volume to produce ice of an appropriate thickness for imaging.

A drawback of the Shake-it-off device is that it tends to produce only small areas of usable ice, with significant variation from grid to grid. Therefore, we sought to develop a robust approach for cryoEM grid preparation with the speed of self-wicking grids, but with the simplicity of conventional grid preparation. We reasoned that the large excess of absorption capacity during conventional grid preparation allows the process to reach the metastable state of a thin film regardless of the exact volume of sample applied to the grid. We hypothesized that if liquid could wick through a grid, a large absorption capacity available on one side of a holey grid would allow specimens to be prepared by high-speed application of small volumes of liquid to the other side (Figure 1*C*). The large absorption capacity would make the technique insensitive to the precise volume of liquid applied, enabling large areas of specimen to be prepared for cryoEM with a simple, robust, and inexpensive droplet generator. We found that this effect could be achieved by treating both surfaces of the grid with extensive glow discharge in air to make them hydrophilic and using glass fiber to wick excess solution through the grid. Following the naming pattern of the Spotiton and Shake-it-off devices, and recognizing the similarity to the backside blotting approach, we name the device presented here *“*Back-it-up*”* or BIU.

## Results and Discussion

### Through-grid wicking with glass fiber allows preparation of thin vitreous ice

We investigated the capability of different blotting materials to rapidly wick liquid through a holey grid when applied to the opposite side of the grid. We found that this effect could be achieved consistently with glass fiber filters but not with conventional filter paper. Glass fiber filters have been used previously to prepare cryoEM specimens from carbon nanotubes in chlorosulfonic acid (Talmon, 2015). Scanning electron microscopy (SEM) with secondary electron detection illustrates the relatively large cellulose fibers in conventional filter paper (Figure 2*A*i) compared to the smaller glass fibers (Figure 2*A*ii). Conventional cryoEM grids with a holey gold film on the rhodium surface of copper-rhodium grids were made hydrophilic by extensive glow discharge in air on both sides and laid with their flat gold surface in contact with either conventional filter paper or a glass fiber filter. A high-speed video of water being applied to the copper surface of the grids shows that the liquid is rapidly wicked through the grid by glass fiber, while this phenomenon did not occur consistently with conventional filter paper (Fig 2*B*; Movie 1).

**Figure 2.**
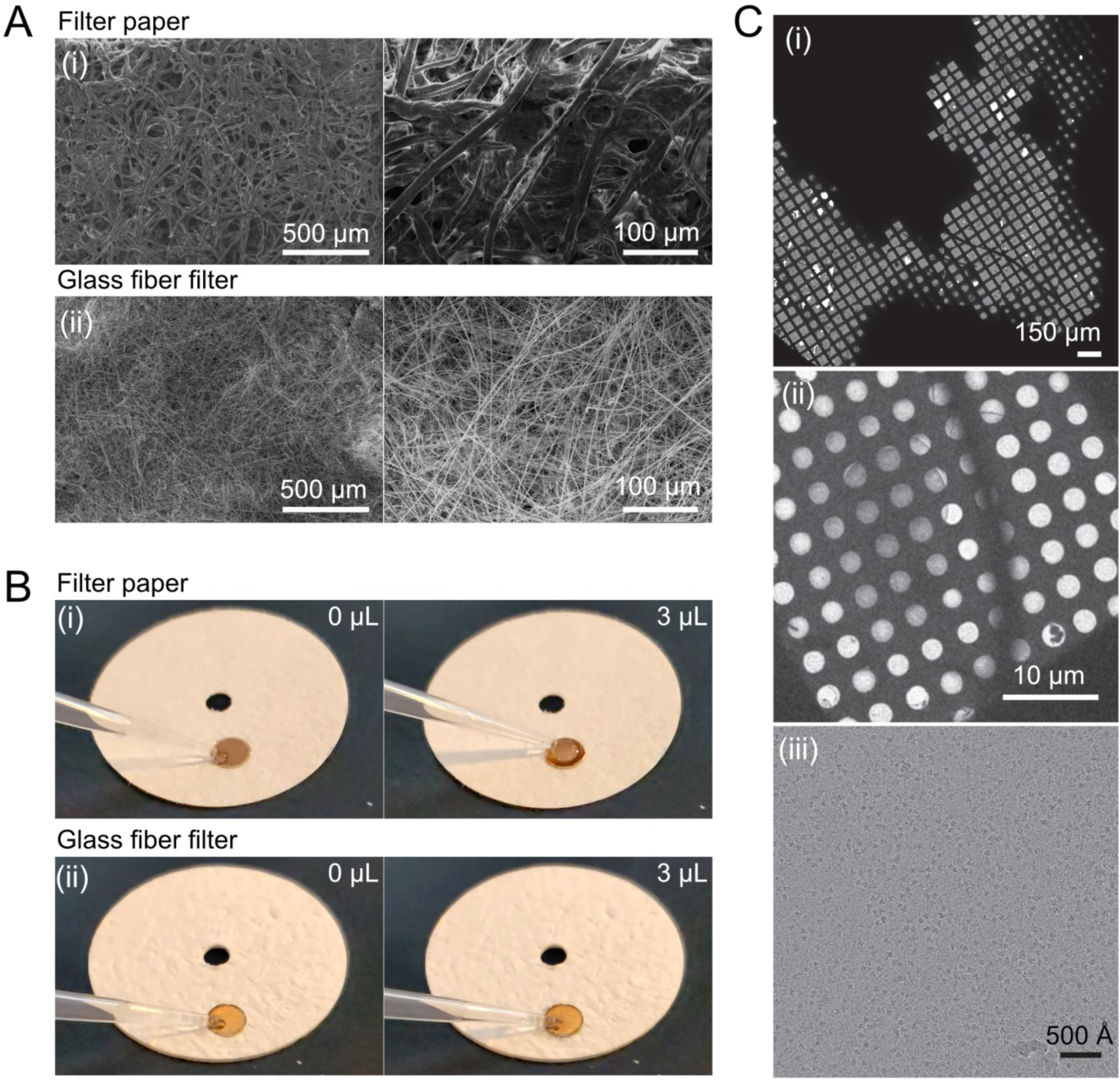
Through-grid wicking with glass fiber. (*A*) Scanning electron microscopy (SEM) images of conventional filter paper (i) and glass fiber filter (ii) at two magnifications. (*B*) A holey gold grid previously subjected to glow discharged on both sides does not consistently show rapid through-grid wicking when in contact with filter paper (i) but water quickly wicks through the grid when it is placed on a glass fiber filter (ii). Movie 1 shows this process in more detail. (*C*) A hemagglutinin trimer specimen prepared by through-grid wicking with a Gatan CP3 grid freezing device and glass fiber filter. A grid atlas (i), grid square (ii), and high magnification image (iii) are shown.

To test if through-grid wicking with glass fiber allows production of thin films of vitreous ice suitable for cryoEM, a specimen of commercially available recombinant hemagglutinin trimer from influenza A virus (Strain A/Hong Kong/1/1968[H3N2]) was prepared with a Gatan Cryoplunge 3 (CP3) grid freezing device. After application of sample to the copper surface of the grid, one sided blotting was performed on the gold surface, with the usual 20 mm circle of filter paper replaced with a 20 mm circle of glass fiber filter. The majority of the 3 µL sample appeared to wick through the grid in ∼1 sec and blotting was continued for a total of 5 sec. A cryoEM atlas of the grid showed a gradient of ice thicknesses (Figure 2*C*i), with many grid squares having ice suitable for data collection (Figure 2*C*ii). High magnification images (Figure 2*C*iii) of holes with appropriate ice for data collection revealed high-contrast particles similar to micrographs obtained previously with the same hemagglutinin specimen (Tan et al., 2017).

### Construction of BIU, a high-speed through-grid wicking specimen preparation device

To develop a device that dispenses small volumes of sample for high-speed cryoEM specimen preparation using the through-grid wicking approach, we adapted the open source Shake-it-off platform (Rubinstein et al., 2019) (Figure 1*C*, 3*A*). In the new instrument design (Figure 3), Dumont L5 tweezers holding an EM grid are attached with a magnetic connector to a large solenoid capable of plunging the grid into a cryogen bath. The cryogen bath, which is insulated by a Styrofoam box, is milled from aluminum and uses liquid nitrogen to maintain a ∼60:40 mixture of propane:ethane in a liquid state (Tivol et al., 2008). The ∼30 mm travel distance of the plunging solenoid when energized is reduced to ∼16 mm by a 3D-printed plastic ring and a magnetic catch keeps the solenoid in its extended position after it is de-energized. A new plastic process mount was designed and 3D-printed to orient the components necessary for through-grid wicking (Figure 3*B*). On one side of the grid, a wicking assembly (Figure 3*C*) uses a tack and a magnet to hold a 20 mm diameter circle of glass fiber filter that is supported by a thin flexible circle of cellulose acetate. This wicking assembly is attached to a small solenoid on the plastic mount so that in its energized position the glass fiber filter is held flush against the grid (Figure 3*D*i). On the other side of the grid the plastic mount holds a 16 mm diameter piezoelectric transducer from an ultrasonic humidifier (Figure 3*E*). The piezoelectric transducer produces a spray of small droplets when a high-frequency signal is applied by the humidifier circuit. A magnet embedded in the Styrofoam insulating box of the cryogen container is held against a reed switch when the box is properly positioned, serving as a safety interlock to prevent premature plunging of the tweezers. The system electronics are controlled, via a custom printed circuit board identical to the one use with the Shake-it-off device, by a Raspberry Pi single board computer running a graphical user interface (GUI) written in *Python3*.

**Figure 3.**
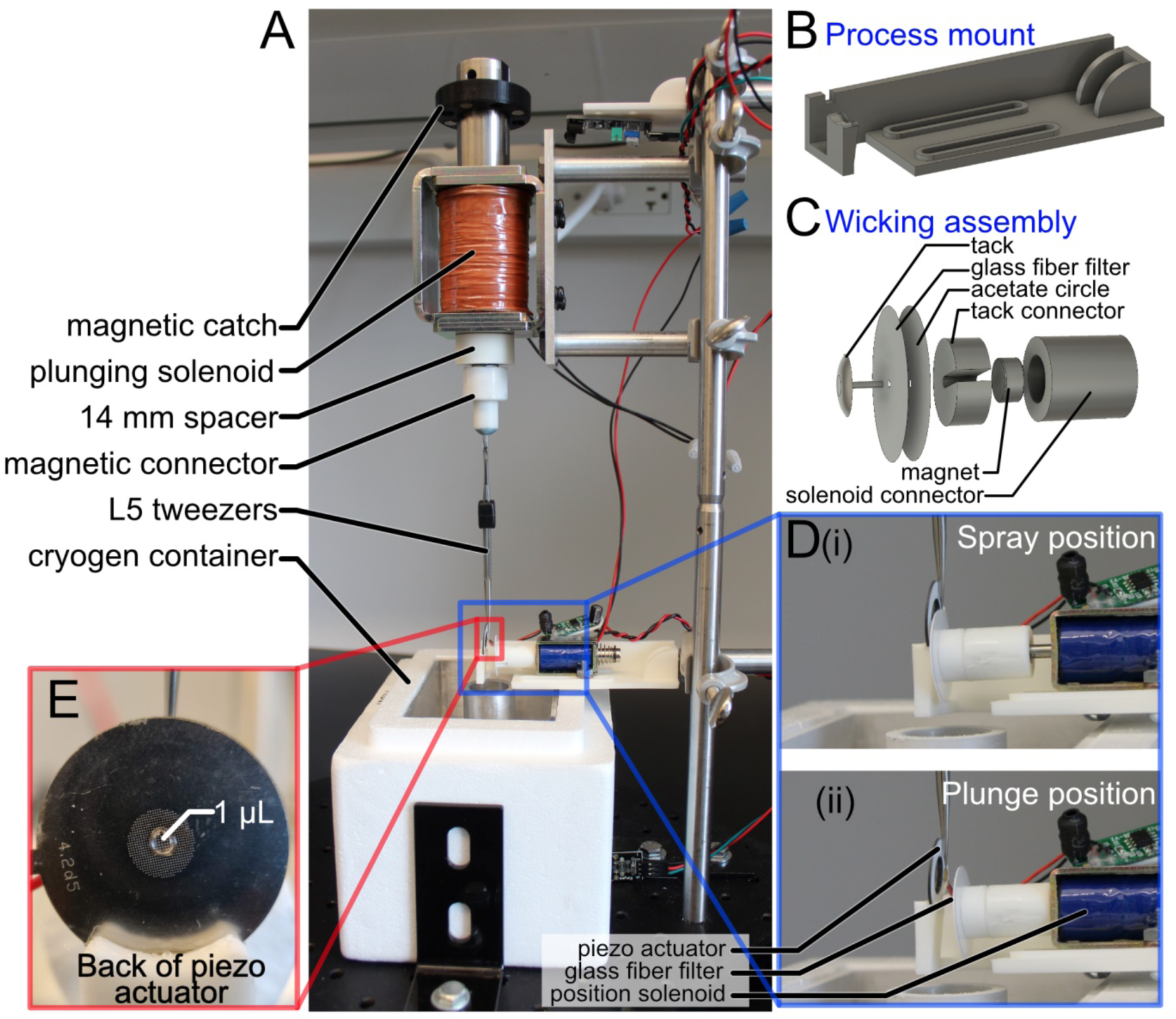
The Back-it-up (BIU) device. (*A*) Fully assembled BIU specimen-preparation device. In comparison with the SIO device, new components are the process mount (*B*) and wicking assembly (*C*). (*D*) When the position solenoid is energized the wicking assembly holds the glass fiber filter flush against the grid to allow spraying and through-grid wicking (i). When the position solenoid is de-energized, the glass fiber filter is retracted creating the clearance needed to plunge the grid into cryogen (ii). (*E*) Sample is applied to the back of the piezoelectric transducer before spraying.

During specimen preparation, a small volume of sample, such as a few hundred nL to 1 µL, is applied to the surface of the piezoelectric transducer opposite the grid. Energizing the high-frequency generating circuit causes a spray of droplets through the piezoelectric transducer onto the copper surface of the grid. After spreading on the grid, liquid is wicked through the grid by the glass fiber filter. Once spraying is complete, the glass fiber filter is held against the grid for an additional wicking time (e.g. 10 msec), after which the glass fiber filter and acetate are pulled back from the grid by de-energizing the small solenoid (Figure 3*D*ii). The large solenoid is then energized and the grid is plunged into the liquid cryogen. At present, this process is carried out at ambient temperature and humidity. However, we see no reason why the BIU device could not be built in a humidity- and temperature-controlled environment, or any number of other controlled environments for specific experiments. Because there is no splash shield for the cryogen, eye protection must always be worn when operating the device. The piezoelectric transducer produces an aerosol from the sample and consequently the device should not be used to prepare samples where aerosols present a health hazard, such as infectious virus particles, some toxins, and infectious prions. However, the device can be installed in a biosafety cabinet to overcome this limitation.

### *High-resolution 3D reconstruction from a specimen prepared with the* BIU *device*

In order to determine if the BIU device is able to prepare specimens amenable to high-resolution cryoEM, we used it to prepare multiple grids with human light chain apoferritin. Spraying was done with the piezoelectric transducer energized for 100 msec, with an additional 100 msec delay after spraying and before plunging. This spraying time is twenty times longer than was used with the Shake-it-off device with self-wicking grids and is sufficient to completely saturate copper-hydroxide nanowire grids, leading to thick ice (Rubinstein et al., 2019). CryoEM atlases of the resulting specimen grids show a gradient of ice thicknesses (Figure 4*A, upper*). This gradient could be due to spatial variation in the amount or droplet size of the spray from the piezoelectric transducer on different parts of the grid. Alternatively, it could be due to the gradient of pressure with which the glass fiber filter is pressed against the grid, with less pressure near the edge of the glass fiber filter circle and more pressure where the glass fiber filter is secured to the wicking assembly by the tack. Nonetheless, grids consistently had approximately one third of their surface covered with ice suitable for imaging. Apoferritin particles can be seen clearly in high-magnification images from multiple grids (Figure 4*A, lower*), demonstrating that the device reproducibly produces specimens with large areas of thin vitreous ice suitable for data collection. A dataset of 3,168 movies was collected from two of the grids (Table 1), allowing calculation of a 3D map at 2.0 Å from 594,259 single particle images (Figure 4*B* and *C*, Figure S1*A* and *B*). An atomic model was built into the density map, including water molecules that could be resolved clearly (Figure 4*C*), confirming that the BIU device is capable of preparing samples that can be used for high-resolution structure determination.

**Table 1.** Cryo-EM data collection and modeling statistics.

|  | Human L Chain<br>Apo ferritin (BIU) | Hemagglutinin<br>(CP3) | Hemagglutinin<br>(BIU) |
| --- | --- | --- | --- |
| **EM data collection / processing** |  |  |  |
| Microscope | FEI Titan Krios |  |  |
| Voltage (kV) | 300 |  |  |
| Camera | Falcon 4 |  |  |
| Mode | Counting |  |  |
| Defocus mean ± std ($\mu\text{m}$) | $0.7 \pm 0.4$ | $2.0 \pm 0.5$ | $2.0 \pm 0.5$ |
| Exposure time (s) | 9 | 9 | 9 |
| Number of fractions | 29 | 29 | 29 |
| Exposure rate ($\text{e}^-/\text{pixel}/\text{s}$) | 3 | 5 | 5 |
| Total exposure ($\text{e}^-/\text{\AA}^2$) | 40 | 40 | 40 |
| Pixel size ($\text{\AA}$) | 0.816 | 1.03 | 1.03 |
| Number of micrographs | 3,168 | 1,284 | 1,556 |
| Number of particles (after cleanup) | 673,794 | 232,841 | 378,143 |
| Number of particles (in final map) | 594,259 | - | 128,305 |
| Symmetry | O | - | C3 |
| Resolution (global) ($\text{\AA}$) | 2.0 | - | 2.9 |
| Local Resolution Range | 1.8-2.2 | - | 2.5-3.4 |
| Directional Resolution Range | 1.9-2.1 | - | 2.7-3.2 |
| Sphericity of 3DFSC | 0.98 | - | 0.98 |
| SCF Value* | 1.00 | - | 0.92 |
| Map B factor ($\text{\AA}^2$) | 92.8 | - | 150.9 |
| **Model statistics** |  |  |  |
| Residue range | 5-176 | - | 9-325, 334-501 |
| Map CC | 0.88 | - | 0.77 |
| RMSD [bonds] ($\text{\AA}$) | 0.006 | - | 0.011 |
| RMSD [angles] ($\text{\AA}$) | 1.003 | - | 1.194 |
| All-atom clashscore | 0.14 | - | 2.45 |
| Average B factors ($\text{\AA}^2$) | | - | |
| Protein | 14.91 | - | 72.39 |
| Ligands | 23.45 | - | 69.53 |
| Water | 16.18 | - | - |
| Ramachandran plot |  | - |  |
| Favored (%) | 98.82 | - | 93.56 |
| Allowed (%) | 1.18 | - | 6.03 |
| Outliers (%) | 0.00 | - | 0.42 |
| Rotamer outliers | 0.00 | - | 0.94 |
| C-β deviations | 0.00 | - | 0.00 |
| MolProbity Score | 0.53 | - | 1.45 |
| EM-Ringer Score | 7.70 | - | 4.07 |

\*The sampling compensation factor (SCF) value is calculated as described (Baldwin and Lyumkis, 2020) and assumes that all orientations have been determined accurately.

**Figure 4.**
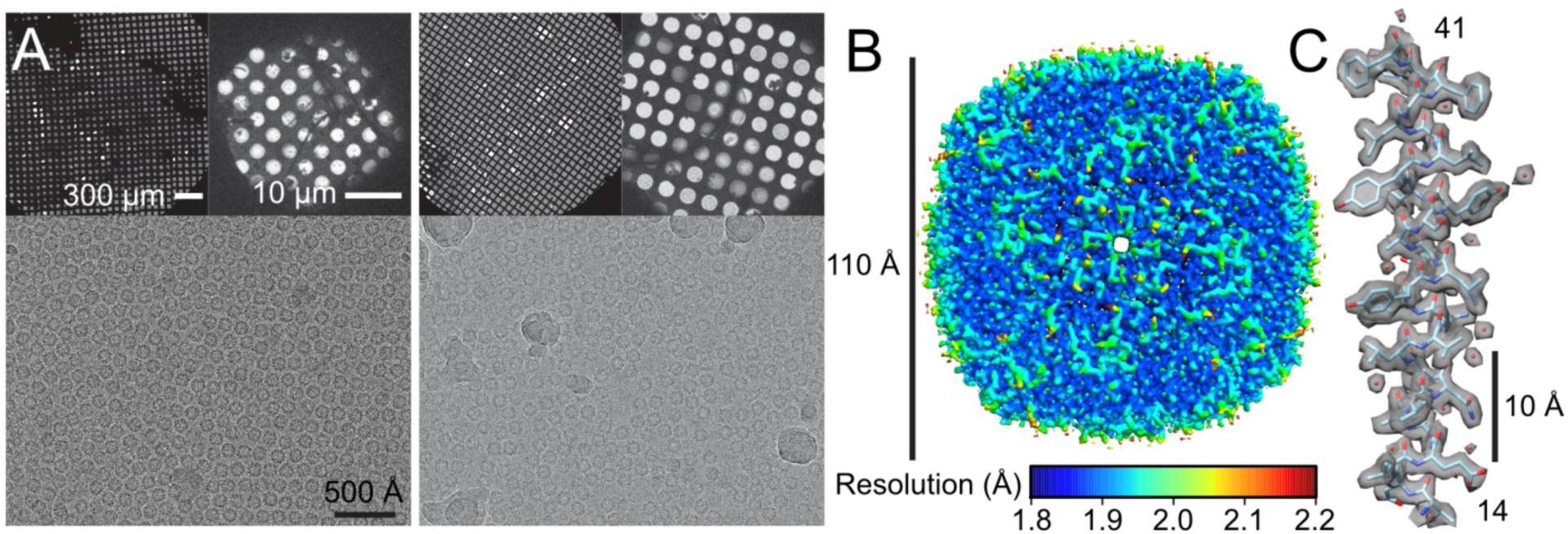
Human apoferritin vitrified using BIU. (*A*) Grid atlases, grid squares, and micrographs from human apoferritin prepared for cryoEM using BIU. (*B*) Local resolution for the apoferritin reconstruction. (*C*) A representative α helix from the apoferritin model (residues 14 to 41) and the experimental density map (grey surface). Water molecules are displayed as red spheres.

### *The* BIU *device can prepare samples of detergent-solubilized membrane proteins*

Membrane proteins maintained in solution with detergents present a unique challenge for cryoEM grid preparation as the detergents often lead to specimens with a steep gradient of ice thickness (Rubinstein, 2007). In order to determine if the BIU device and the through-grid wicking approach can prepare high-quality cryoEM specimens when detergents are present in samples, we vitrified cryoEM grids of two different membrane proteins samples that included *n*-dodecyl β-D-maltoside (DDM). These samples were a yeast ATP synthase (Guo et al., 2017) (Figure 5*A*) and a bacterial ATP synthase complex (Figure 5*B*). Similar to the apoferritin grids, large areas of usable ice were produced for these specimens. High-magnification images from thin vitreous ice show the membrane protein complexes clearly. These particle images allowed calculation of 2D class average images with high-resolution features, confirming that the BIU device can be used with detergent-containing membrane protein samples.

**Figure 5.**
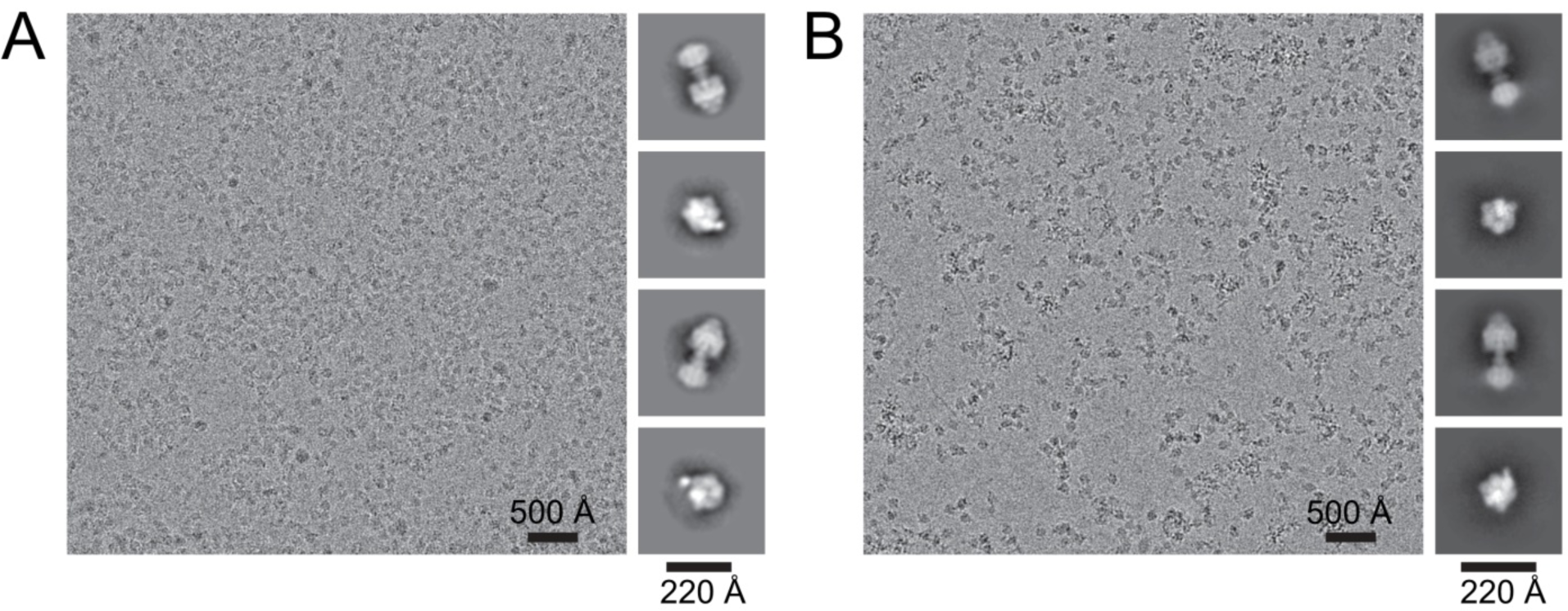
Membrane proteins solubilized in detergent vitrified using BIU. (*A*) Representative high-magnification image of yeast ATP synthase and r2D class averages. (*B*) Representative high-magnification image of bacterial ATP synthase complex and representative 2D class averages.

### High-speed through-grid wicking can reduce time-dependent preferred orientations

With the Spotiton device and self-wicking grids, the influenza A hemagglutinin trimer was shown to undergo time-dependent adoption of a preferred top-view orientation at the air-water interface during grid freezing (Noble et al., 2018). This preferred orientation can prevent calculation of a 3D map. In that study, conventional use of a Gatan CP3 (∼1 sec blot time) resulted in most particles adopting the three-fold symmetric top views and only 4% of particle images assigned to classes that show a side view of the trimer. In contrast, the number of particle images contributing to side view classes increased to 9% with the Spotiton device and a 500 msec spot-to-plunge time, and to 19% with a 100 msec spot-to-plunge time (Noble et al., 2018).

To determine if specimen preparation with through-grid wicking can be performed at sufficient speed to reduce adoption of preferred orientations, grids were prepared with the BIU device using the identical recombinant hemagglutinin trimer. Sample was sprayed onto the grid with the piezoelectric transducer energized for 30 msec, with an additional 30 msec delay after spraying and before plunging to allow wicking of excess solution by the glass fiber filter. Recording this process at 480 fps (frames per second) showed that the piezoelectric circuit was active for ∼45 msec, followed by an additional ∼30 msec where the grid remained in contact with the glass fiber filter, and ∼60 msec passing during which the glass fiber filter was fully retracted and the grid plunged into cryogen (Figure 6*A*, Movie 2). This sequence constitutes an overall process time of 135 msec, with immersion in cryogen occurring 90 msec after the end of spraying and 60 msec after breaking contact between the grid and glass fiber filter. 1,556 movies were acquired from this specimen. As with the apoferritin and ATP synthase specimens, approximately one third of the grid had ice that was suitable for imaging. 1,284 movies were also acquired from a sample of the hemagglutinin trimer, described earlier, that was prepared using the Gatan CP3 with glass fiber filter paper and backside blotting. With the Gatan CP3 and a 5 sec blot time, 2% of particle images (1,506 out of 91,879) were assigned to a single class corresponding to a side view, while 98% (90,373 out of 91,879) were assigned to classes that showed top views or oblique views (Figure 6*B*). In contrast, with the BIU grid 66% of particle images (170,419 out of 256,303) were assigned to top or oblique view classes while 34% (85,884 out of 256,303) were assigned to side-view classes (Figure 6*B*). The top and oblique views for the BIU specimen also appeared to include more tilted oblique views than for the Gatan CP3 specimen (Figure 6*B, left*). The large number of single particle images presenting side views in the BIU specimen allowed 3D reconstruction of the hemagglutinin trimer at 2.9 Å resolution (Figure 6*C*, Figure S2*A* and *B*, Table 1) without the need for tilting the specimen (Tan et al., 2017). The high quality of the map allowed construction of an atomic model, including three N-linked glycans (Figure 6*D*, *bottom right*). The map has a sampling compensation factor (Baldwin and Lyumkis, 2020) of 0.92 and a 3D Fourier shell correlation sphericity (Tan et al., 2017) of nearly 1.00 when calculated either between half-maps or between the map and the model (Figure 6*C*, Figure S2*B*, Table 1). These statistics support the directional isotropy of the resolution. Local resolution in the map is slightly worse near the membrane proximal part of the trimer (Figure 6*C, bottom of map*) as was suggested previously for this complex (Vilas et al., 2020), likely due flexibility in the structure.

**Figure 6.**
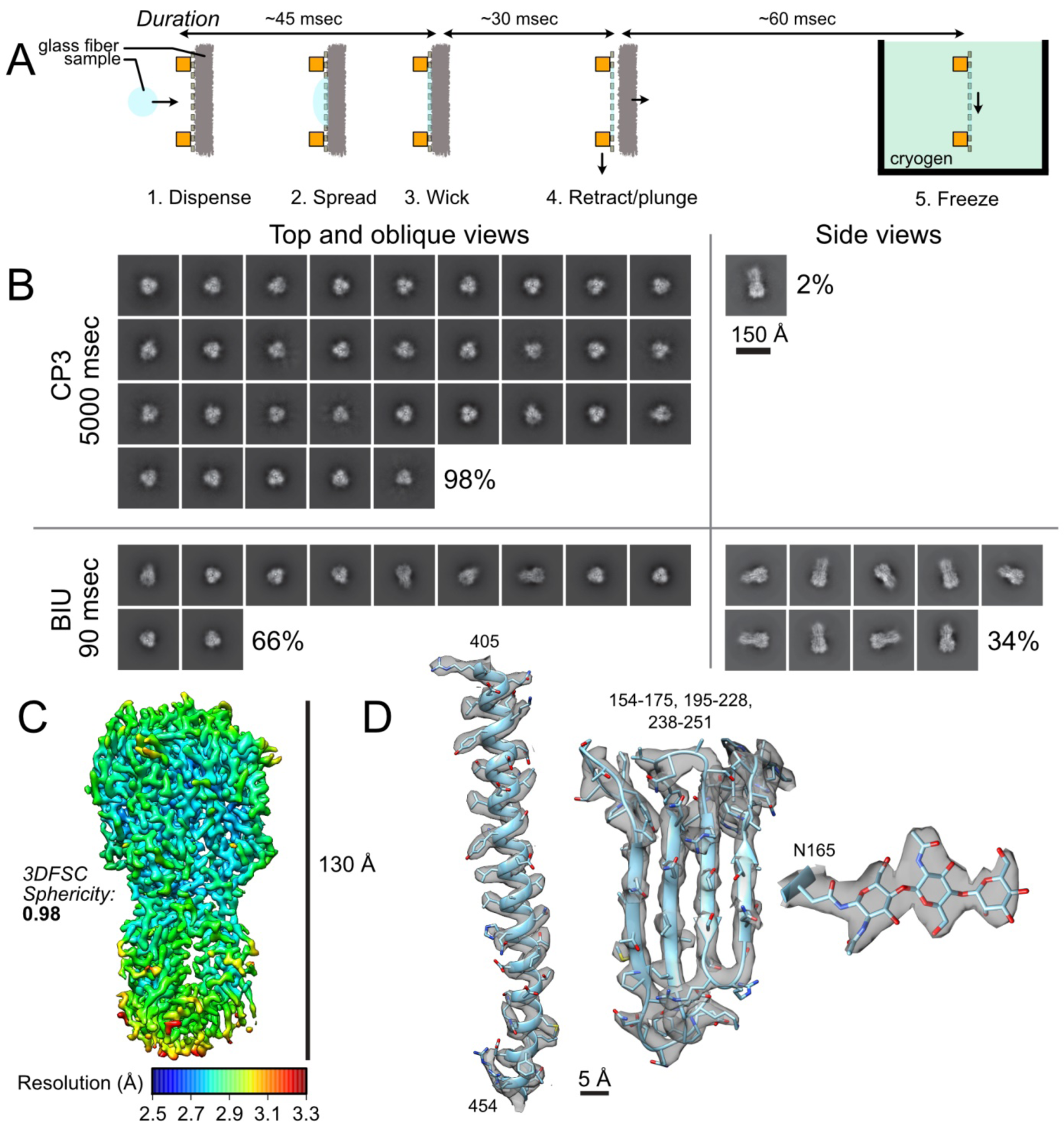
High-speed freezing with BIU improves the orientation distribution in a hemagglutinin trimer specimen. (*A*) Timing of the grid freezing process measured with high-speed video. (*B*) The complete set of 2D class averages from hemagglutinin trimer specimens prepared with the Gatan CP3 and BIU. The fraction of particle images contributing to top and oblique view classes versus side view classes is indicated. (*C*) Local resolution map of the hemagglutinin trimer. (*D*) A representative α helix (residues 405-454), β sheet (residues 154-175, 195-228, 238-251), and N-glycan (residue 165) from the hemagglutinin model, overlaid with the experimental density map (grey surface).

## Discussion

Implementation of through-grid wicking for high-speed cryoEM specimen preparation allows grids to be frozen in tens of milliseconds, rather than the multiple seconds needed for conventional grid preparation. The method does not require the self-wicking nanowire grids used previously in the Spotiton, Chameleon, and Shake-it-off devices. Avoiding nanowire grids and using conventional holey cryoEM grids simplifies the consumables needed for grid preparation and reduces a source of variability that has affected specimen preparation with nanowire grids (Wei et al., 2018). By providing a high capacity for absorbing liquid, through-grid wicking appears to increase the tolerance for excess-liquid applied to the grid. This decrease in sensitivity avoids creation of large areas of ice that are too thick for data collection, even when approximately six to twenty times the spraying time of the Shake-it-off device (Rubinstein et al., 2019) is used with the BIU device, or 3 µL of sample is applied directly to the grid in the Gatan CP3. As a result, large areas of high-quality ice can be prepared readily and reproducibly with the BIU device, with approximately one third of the grid surface available for data collection. This large usable area allows for straightforward structure determination from a single grid.

Electron tomography of specimens prepared with the Spotiton device showed that the mechanism for reducing preferred orientations was decreased interaction with the thin film air-water interface before freezing (Noble et al., 2018). Although repeating the mechanistic analysis of this effect with the BIU device is beyond the scope of the present manuscript, which instead focuses on the ability of the approach to prepare usable specimens, the abundance of hemagglutinin trimer side views is presumably due to the same effect that occurs when performing high-speed grid preparation with self-wicking grids.

In the experiments described above we used the same ultrasonic specimen application approach employed with the Shake-it-off instrument, which is related to an acoustic wave atomization approach described previously (Ashtiani et al., 2018). However, through-grid wicking can almost certainly be used with other methods for applying sample onto an EM grid. The piezoelectric transducer from an ultrasonic humidifier used here has the advantage of having no ‘dead volume’ of sample that must be loaded into the device but cannot be dispensed. It has the disadvantage of using relatively large volumes for each grid, such as hundreds of nanoliters up to one microliter, although it may be possible to reduce this volume further. Alternative sample application methods include the ink-jet approach implemented in the Spotiton device (Dandey et al., 2018; Jain et al., 2012) and the microfluidic sprayers used with several time-resolved specimen preparation devices (Feng et al., 2017; Kaledhonkar et al., 2018; Kontziampasis et al., 2019; Lu et al., 2014). Through-grid wicking could also prove useful for preparing optimal ice when applying nanoliter samples to a grid with a microcapillary (Arnold et al., 2017, 2016) or from a solid surface (Ravelli et al., 2020).

Similar to the Shake-it-off platform on which it is based, the BIU instrument is inexpensive to construct. The implementation described here cost ∼C$1,000 (∼USD$700) to build, or approximately 500 to 1000× less than a commercially available second-generation grid preparation device. All components needed to build the device are either commercially available or can be made by online 3D printing, printed circuit board fabrication, and CNC milling services, so that specialized manufacturing infrastructure is not required. The 3D design files, the printed circuit board design files, and the control software are available online to allow others to easily replicate the instrument (https://github.com/johnrubinstein). We believe that the reproducibility and large areas of usable ice from through-grid wicking, combined with the simplicity and low construction cost of the BIU device, provides a robust and easily accessible approach to high-speed cryoEM grid preparation for a variety of specimens.

## Author contributions

JLR devised the grid preparation approach, built the instrument, and wrote the control software. YZT devised the characterization strategy, prepared samples, collected data, performed image analysis, and constructed atomic models. JLR and YZT wrote the manuscript and prepared the figures

## Acknowledgements

Recombinant human apoferritin was a gift from Taylor Sicard and Jean-Philippe Julien (The Hospital for Sick Children). Recombinant yeast ATP synthase was prepared by Hui Guo (The Hospital for Sick Children). Recombinant bacterial ATP synthase was prepared by Gautier Courbon (The Hospital for Sick Children). We thank Ali Darbandi (The Hospital for Sick Children) for performing the SEM imaging in Figure 2. We thank Drs. Glaeser and Han (Lawrence Berkeley National Laboratory) for confirming that the through-grid wicking phenomenon is not due to surfactant on the glass fiber filter. YZT was supported by a postdoctoral fellowship from the Canadian Institutes of Health Research and JLR was supported by the Canada Research Chairs program. This work was supported by a Discovery Grant from the Natural Sciences and Engineering Research Council. CryoEM data was collected at the Toronto High-Resolution High-Throughput cryoEM facility, supported by the Canada Foundation for Innovation and Ontario Research Fund.

## Data availability

All raw movies, micrographs, particle stacks, and relevant metadata files will be deposited into EMPIAR. CryoEM maps of apoferritin and hemagglutinin have been deposited into the EMDB as EMD-21951 and EMD-21954 respectively. Models of apoferritin and hemagglutinin have been deposited into the PDB as 6WX6 and 6WXB. Original software and 3D design files are available from https://github.com/johnrubinstein

---

**Figure S1.**
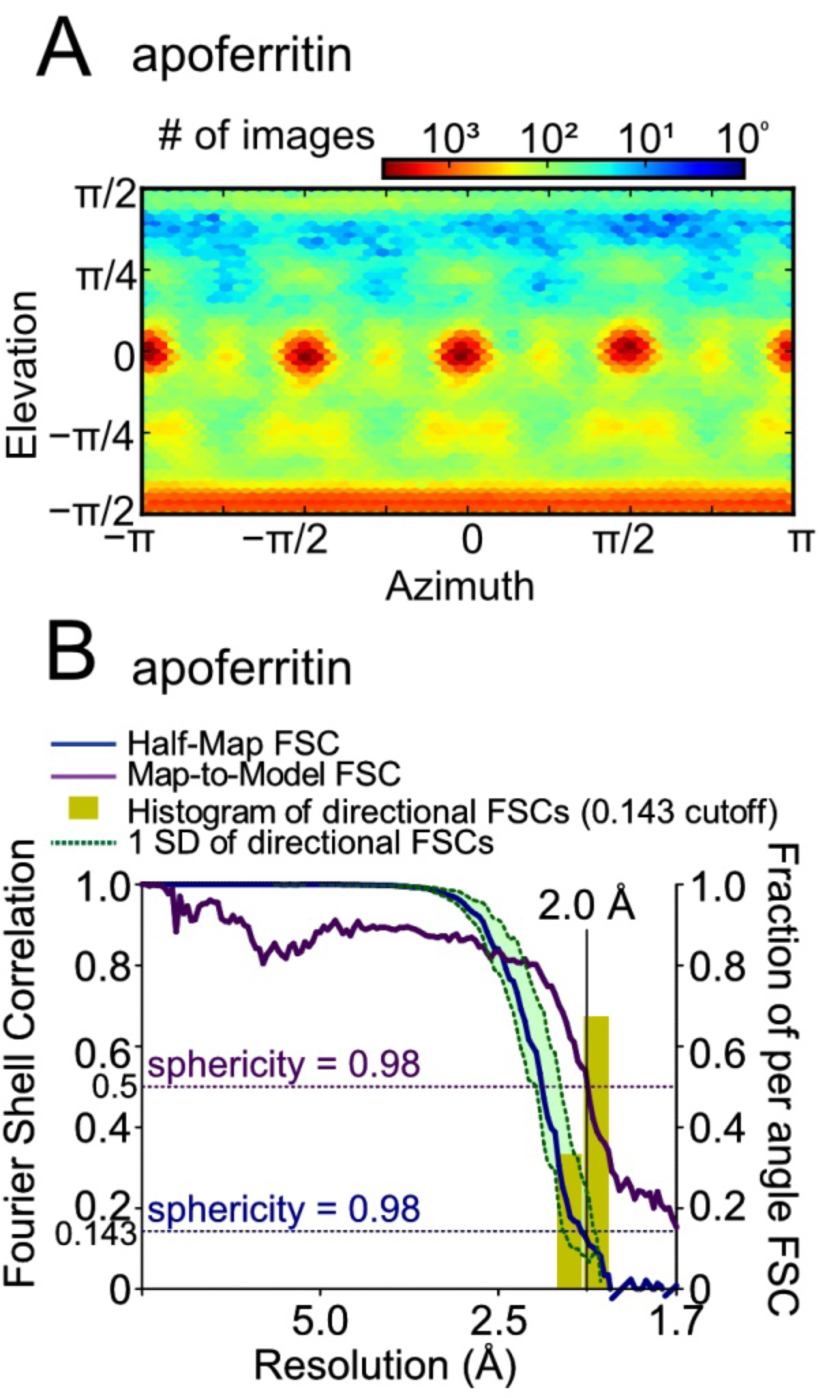
CryoEM map validation. (*A*) Euler angle distribution for human light chain apoferritin. *(B)* Fourier shell correlation (FSC) curves for human light chain apoferritin show the map resolution from half-map (blue) and map-to-model (purple) correlation at FSC=0.143 and FSC=0.5, respectively. A histogram of directional resolutions sampled evenly over the 3DFSC (yellow) is also shown and the corresponding sphericity is indicated.

**Figure S2.**
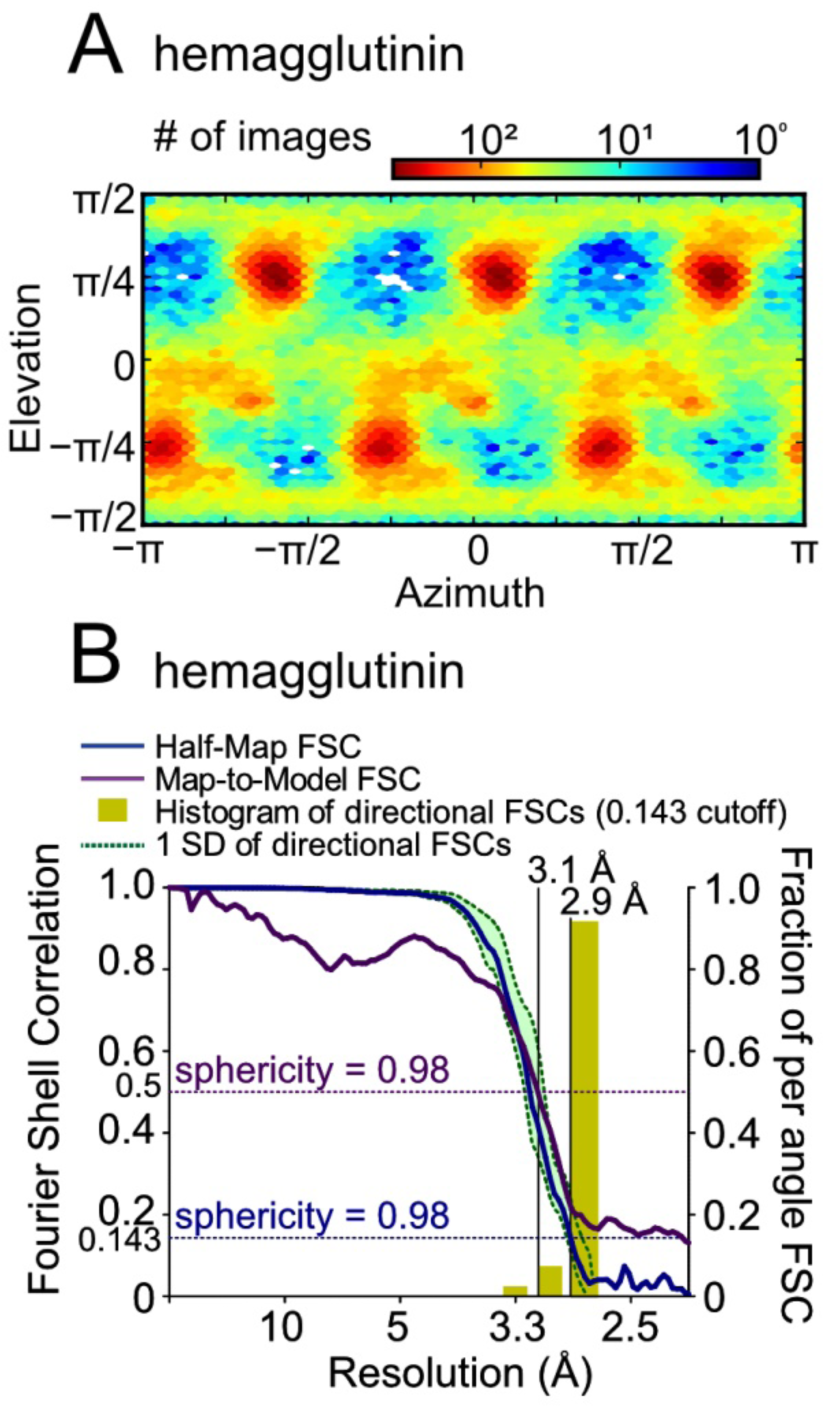
CryoEM map validation. (*A*) Euler angle distribution for influenza A H3N2 hemagglutinin trimer. *(B)* Fourier shell correlation (FSC) curves for influenza A H3N2 hemagglutinin trimer show the map resolution from half-map (blue) and map-to-model (purple) correlation at FSC=0.143 and FSC=0.5, respectively. A histogram of directional resolutions sampled evenly over the 3DFSC (yellow) is also shown and the corresponding sphericity is indicated.

**Movie 1. Comparison of filter paper and glass fiber filter for through-grid wicking.** Water (5 µL) applied to a hydrophilic holey gold grid does not consistently wick through the grid when it is placed on filter paper (top) but quickly wicks through the grid when it is placed on a glass fiber filter (bottom).

**Movie 2. High speed video of the BIU process.** A video recorded at 480 frames per second shows the high-frequency generating circuit of the piezoelectric transducer energized for ∼45 msec. A further 30 msec elapses where the glass fiber filter remains in contact with the grid. The grid is plunged into cryogen ∼60 msec after it breaks contact with the glass fiber filter.

## Materials and Methods

### Device design, construction, and consumables

New 3D parts were designed with *Autodesk Fusion 360* and printed in polylactic acid with a Raise3D Pro2 3D printer. All other parts and components were identical to those used with the SIO device (Rubinstein et al., 2019). 20 mm circles of glass fiber filters (VWR grade 691) and filter paper (Whatman #1) were cut with the circle punch from the Gatan CP3 grid freezing device. Nanofabricated holey gold film coated EM grids with regularly spaced ∼2 µm holes were prepared on EMS Maxtaform 400 mesh Cu/Rh grids as described previously (Marr et al., 2014). Grids were glow discharged in air for two minutes on each side with a PELCO easiGlow (Ted Pella) immediately before use.

### Apoferritin specimen preparation

Human apoferritin light chain (UniProtKB: P02792), modified with an N-terminal Strep tag and linker, a C-terminal extension, and a K173R mutation, was a gift from Taylor Siccard and Jean-Philippe Julien (The Hospital for Sick Children). Apoferritin was diluted to 5 to 10 mg/mL for specimen preparation with BIU. Sample (1 µL) was applied to the piezoelectric transducer and liquid was transferred to the grid by energizing the high-frequency generating circuit for 100 msec. An additional 100 msec was allowed for the glass fiber filter to wick sample through the grid before retracting the filter and plunging the grid into the liquid propane/ethane bath.

### Yeast ATP synthase specimen preparation

Yeast ATP synthase was a gift from Hui Guo. Yeast ATP synthase was diluted to 10 mg/mL with 0.05% (w/v) DDM for specimen preparation with BIU. Sample (1 µL) was applied to the piezoelectric transducer and liquid was transferred to the grid by energizing the high-frequency generating circuit for 50 msec. An additional 1000 msec was allowed for the glass fiber filter to wick sample through the grid before retracting the filter and plunging the grid into the liquid propane/ethane bath.

### Bacterial ATP synthase specimen preparation

Bacterial ATP synthase was a gift from Gautier Courbon. The protein was diluted to 2 mg/mL with 0.05% (w/v) DDM for specimen preparation with BIU. Sample (1 µL) was applied to the piezoelectric transducer and liquid was transferred to the grid by energizing the high-frequency generating circuit for 10 msec. An additional 500 msec was allowed for the glass fiber filter to wick sample through the grid before retracting the filter and plunging the grid into the liquid propane/ethane bath.

### Hemagglutinin specimen preparation

Hemagglutinin (MyBioSource product MBS434205) was used without dilution (1 mg/mL). For specimen preparation with the Gatan Cryoplunge 3, sample (3 μL) was applied onto the copper side of nanofabricated gold grids in the environmental chamber of the device (>80% RH, 298 K). Grids were blotted from only the gold side of the grid with glass fiber filter for 5 sec and plunged into liquid ethane. For freezing hemagglutinin with the BIU device, sample (1 µL) was applied to the piezoelectric transducer and liquid was transferred to the grid by energizing the high-frequency generating circuit for 30 msec. An additional 30 msec was allowed for the glass fiber filter to wick sample through the grid before retracting the filter and plunging the grid into the liquid propane/ethane bath. High-speed movies were recorded at 480 frames per second with a rolling shutter using a OnePlus6 mobile phone.

### CryoEM data collection

ATP synthase data was acquired at 200 kV with a FEI Tecnai F20 electron microscope and Gatan K2 summit camera operating in electron counting mode. Data collection was done manually using Digital Micrograph software. 50 movies were collected for the yeast ATP synthase, and 107 movies were collected for the bacterial ATP synthase at a pixel size of 1.45 Å/pixel. Movies were collected for 15 sec with 30 exposure fractions, a camera exposure rate of ∼8 e^-^/pix/sec, and total specimen exposure of ∼50 e^-^/Å^2^.

For apoferritin and hemagglutinin, data was acquired at 300 kV with a Thermo Fisher Scientific Titan Krios G3 electron microscope and prototype Falcon 4 camera operating in electron counting mode at 250 frames/sec. The microscope was automated with the *EPU* software package and data collection was monitored with *cryoSPARC Live* (Punjani et al., 2017).

3,168 movies were acquired from two grids of apoferritin prepared with BIU. The pixel size was calibrated at 0.816 Å/pixel by fitting a crystal structure of human ferritin L chain (PDB ID: 2FFX) (Wang et al., 2006) into a preliminary 3D map. Movies were collected for 9 sec with 29 exposure fractions, a camera exposure rate of ∼3 e^-^/pix/sec, and total specimen exposure of ∼40 e^-^/Å^2^. A 100 μm objective aperture was used. 1,284 movies were acquired from one grid with hemagglutinin prepared with the CP3. The pixel size was calibrated at 1.03 Å/pixel from a gold diffraction standard. Movies were collected for 9 sec with 29 exposure fractions, a camera exposure rate of ∼5 e^-^/pix/sec, and a total specimen exposure of ∼40 e^-^/Å^2^. No objective aperture was used. 1,556 movies were acquired from one grid with hemagglutinin prepared with the BIU device. The calibrated pixel size was 1.03 Å. Movies were collected for 9 sec with 29 exposure fractions, a camera exposure rate of ∼5 e^−^/pix/sec, and a total specimen exposure of ∼40 e^-^/Å^2^. No objective aperture was used.

### Apoferritin image processing

Exposure fractions were aligned with *MotionCor2* (Zheng et al., 2017) within *Relion* (Kimanius et al., 2016; Scheres, 2012), using 7×7 patches and a B factor of 500 pixel^2^ for global alignment and 150 pixel^2^ for local alignment. Micrographs were then imported into *cryoSPARC* (Punjani et al., 2017) and patch CTF estimation was performed. Templates generated from 2D classification during the *cryoSPARC Live* session were used for template selection of particles. 2D classification was used to remove junk particles, resulting in a dataset of 673,794 particle images. An additional round of 2D and 3D classification reduced the dataset to 594,259 particle images. Homogeneous refinement with global and local CTF refinement was then performed and the refined defocus values, Euler angles, and shifts were exported to *Relion*. Particle polishing was done in *Relion* (Zivanov et al., 2019). The particle images were imported back into *cryoSPARC* for a final round of homogeneous refinement with global and local CTF refinement, producing a 2.0 Å map. Octahedral symmetry was enforced throughout all 3D refinement steps. Transfer of data between *Relion* and *cryoSPARC* were done with *pyem* (<u>10.5281/zenodo.3576630</u>).

### Hemagglutinin image processing

Movies were aligned with patch motion correction and patch CTF estimation was performed in *cryoSPARC*. Particle images were selected by blob picking with multiple rounds of 2D classification to remove junk particles. This process resulted in 232,841 hemagglutinin particle images for the specimen prepared with the CP3 grid freezing device. The process gave 378,143 hemagglutinin particle images for the specimen prepared with the BIU device. For 3D structure determination, local motion correction was performed on these particles, followed by repeated rounds of *ab initio* 3D classification with two classes to remove junk particles. The final stack of 128,305 particle images was used to produce a map at 2.9 Å resolution by homogeneous refinement with C3 symmetry enforced and CTF refined globally and locally.

### Model building and refinement

A crystal structure of human light chain ferritin (PDB ID: 2FFX) (Wang et al., 2006) was used to start model building for apoferritin and a crystal structure of the influenza A H3N2 hemagglutinin (PDB ID: 3WHE) (Iba et al., 2014) was used to start model building for hemagglutinin. Models were built manually in *Coot* (Emsley and Cowtan, 2004) and refined with iterative rounds of adjustment in *Coot* and real space refinement with *Phenix* (Adams et al., 2010; Afonine et al., 2018). This process was continued until statistics calculated with *Molprobity* (Chen et al., 2010) ceased to improve. Restraints for the ligands were generated with *phenix.ready_set*. The calcium ion in the apoferritin structure was identified by comparison with a previous apoferritin structure (PDB ID: 1Z6O) (Hamburger et al., 2005). The final map and model were validated using *EMRinger* (Barad et al., 2015) to compare map to model, *cryoSPARC*’s implementation of *blocres* (Cardone et al., 2013) to assess map local resolution, the *3DFSC* suite (Tan et al., 2017) to calculate directional resolution anisotropy, and the *SCF* program (Baldwin and Lyumkis, 2020) to calculate the sampling compensation factor (SCF), which quantifies how preferred particle orientations contribute to attenuation of the Fourier shell correlation (FSC). Map-to-model FSCs were calculated by converting the model to a map with the *molmap* function in *Chimera* at the Nyquist resolution. A mask was made from this map with *Relion* (after low-pass filtering to 8 Å, extending by 1 pixel, and applying a cosine-edge of 3 pixels), and was applied to the map. Map-to-model FSC was calculated with *proc3d* in *EMAN* (Ludtke et al., 1999).

